# Interactions between the representations of pain and reward suggest dynamic shifts in reference point

**DOI:** 10.1101/2023.07.20.549309

**Authors:** R Hoskin, C Pernet, D Talmi

**Author notes:** Corresponding author: Deborah Talmi Department of Psychology, University of Cambridge, Downing Site, Cambridge.

## Abstract

With an aim to understand how brains compute the expected utility of mixed prospects, namely those associated with both negative and positive attributes, we designed a task which equated the opportunity to learn about these attributes and their hedonic value. Participants underwent fMRI scanning while they experienced a classical conditioning paradigm where emotionally-neutral faces predicted a probability of pain and reward conforming to a 2 (Electric Pain: high, low) x 2 (Monetary Reward: high, low) factorial design. We found a robust interaction between the anticipation of pain and reward in the BOLD signal. Analysis of simple effects revealed that sensitivity to each attribute increased under high levels of the other attribute. In the bilateral insula and mid-cingulate gyrus sensitivity to pain was greater under high reward, and in the OFC, caudate, ventral striatum and VTA sensitivity to reward was greater under high pain. We speculate that this pattern is due to dynamic shifts in the reference point participants considered to evaluate each attribute.

## Introduction

Going for drinks with colleagues is fun, but leaving early means that work piles up for the following day. This outing thus has two very different consequences, one with a positive subjective value, and one with a negative one. Although many prospects have attributes that are so different that they appear ‘incommensurable’, they can nevertheless be compared through their subjective utility (Chib et al., 2009; FitzGerald et al., 2009; Levy & Glimcher, 2012). Prospect theory (Kahneman & Tversky, 1979), a dominant economic theory in neuroeconomics research, considers that the subjective utility of prospects that have both positive and negative attributes is computed by subtracting the (dis)utility of the negative attribute from the utility of the positive attribute. This economic framework has been applied to our understanding of complex interactions between positive and negative prospect attributes, even when they are not monetary. For example, it has been used to explain situations where extreme motivation to achieve a goal leads to insensitivity to pain, such as when injured persons appear oblivious to pain while they act to increase the survival chances of themselves and others (Leknes & Tracey, 2008). In neuroscience experiments, prospect theory is used routinely to interrogate how participants integrate the cost and value of goods (e.g. Hare et al., 2008). The economic framework links well with the behaviourist conceptualisation of appetitive and aversive processes as opponent (Konorski, 1967)), which had important influence on neuroeconomics research (Seymour et al., 2005). It also connects well with the conceptualisation, in psychology, of emotional valence as a latent variable represented along a bipolar continuum that ranges from ‘unpleasant’ to ‘pleasant’ (Russell & Barrett, 1999), possibly represented in the orbitofrontal cortex (Chikazoe et al., 2014).

Yet there are also alternative ways to think about the interaction between pain and reward. Psychologists have sometimes described dimensions of positive valence and negative valence as two independent, potentially coactive, dimensions (Cacioppo & Berntson, 1994). This understanding aligns with the failure to find a robust representation of bipolar valence in human neuroimaging studies (Lindquist et al., 2015), which contrasts with the far easier task of pinpointing a neural expression of the valence of emotional arousal in the same meta-analysis, and with the robust evidence in neuroimaging experiments for a representation of positive valence (Lindquist et al., 2015) and reward (Bartra et al., 2013) in the ventromedial PFC. There is also evidence that even when the same brain regions are involved in processing positive and negative stimulus attributes, the sub-regions of such representations may be separable, both in animals and in humans (Berridge & Kringelbach, 2015; Seymour et al., 2007). Intriguingly, evidence that negative and positive prospect attributes interact, for example when depression decreases reward sensitivity (Alloy et al., 2016; Huys et al., 2013), anhedonia increases punishment sensitivity (Ossola et al., 2023) or the threat of pain increases reward sensitivity in healthy controls (Park et al., 2011; Talmi et al., 2009), suggests that at some level, they must be represented separably. This argument ties with increasing evidence that humans represent both dimensions such as valence and arousal, and discrete emotions (Decoding the Nature of Emotion in the Brain, 2016). Taken together, it is not fully clear how the brain evaluates “mixed prospects”, namely, outcomes which have at least two incommensurable attributes with opposite valence (Talmi & Pine, 2012).

To advance understanding of multi-attribute prospects, researchers have paired monetary and a non-monetary outcomes. For example, the value of a good such as a household appliance, together with monetary wins and losses, can be computed by translating the value of the good to monetary units (FitzGerald et al., 2009; Hare et al., 2008). Three neuroimaging studies examined mixed prospects, where the second, non-monetary attribute was negative. Two studies used physical pain, and reported that the representation of either the money outcome or the pain outcome was weaker when it was accompanied by an outcome with the opposite sign. Although the studies used different tasks, both administered positive and negative stimuli in a factorial design, with positive valence operationalized through monetary gains, and negative valence through electric pain stimulation (Bulganin et al., 2014; Choi et al., 2013). Bulganin and colleagues first paired neutral stimuli with pain (CS+) or safety (CS-). They then examined hemodynamic responses during a counterconditioning phase, where the CS+ was paired with reward, in the absence of pain, and then re-presented the USS and examined responses to the CSs during extinction. Choi and colleagues examined responses during the delay that followed the presentation of pain-predicting CSs, but before the simultaneous onset of both the pain US and an instrumental task, where participants could win reward. Both studies reported an interaction between the positive (monetary) and negative (pain) outcomes in the Ventral tegmental area (VTA) and striatum, such that the response to reward in these areas decreased when the threat of a negative pain outcome increased, and the response to the anticipation of pain decreased when it was to be potentially accompanied by a higher reward. The amygdala also expressed differential effects of reward as a function of pain in Bulganin et al.’s study, but only the main effect of reward in Choi et al.’s study. These findings accord with the interpretation that the VTA and striatum represent a correlate of subjective utility (Levy & Glimcher, 2012). The third study (Yee et al., 2021) paired one of three monetary outcomes (small, medium and large) and the delivery of a one of three liquids with positive, negative, or neutral valence. The authors found that only the dorsal cingulate cortex (but not the striatum, ventromedial PFC or anterior insula) was sensitive to both prospect attributes, and that activity there corresponded to the subjective utility of the combined outcomes.

Although these studies help us understand where incommensurable prospect attributes may be integrated, the differences between the ways participants learned about the positive and negative attributes complicate the interpretation of the results. For example, in both studies the representation of pain preceded the establishment of reward representation. The counterconditioning manipulation in Bulganin et al.’s study involved introducing the reward in combination with threat extinction (i.e. the removal of the previously established pain conditioning). In Choi et al.’s study participants received the pain while engaged with an instrumental task facilitating rewards. In all three studies the uncertainty of pain was not the same as the uncertainty of reward. In Bulganin et al.’s study uncertainty about pain emerged gradually during the counter-conditioning phase, and extinction took place immediately after trials where CSs and reward were paired, but a lot later after the last trials where CSs and pain were paired, so uncertainty about reward delivery may have been higher than uncertainty about pain delivery. In Choi et al.’s and Yee et al.’s studies only reward was dependent on participants’ behaviour, and was in that sense both less certain, and more controllable, than the negative outcome, which was classically conditioned in the former and delivered randomly in the latter study.

The relationship between the neural representations of positive and negative feelings, and between the neural representation of reward and punishment, is of continued interest, evident in meta-analyses devoted to these issues (Bartra et al., 2013; Lindquist et al., 2015); but meta-analyses cannot shed light on the way these representations interact online. To move towards a fuller understanding of how brains compute the expected utility of mixed prospects we designed a task where the manipulation of each attribute was formally and hedonically equivalent. To eliminate the influence of expected utility on control, evident in a previous study (Yee et al., 2019), we employed a simple design without an element of choice, based on Pavlovian conditioning (following Sescousse et al., 2013). Participants performed a Pavlovian conditioning task conforming to a 2 (Electric Pain: high, low) x 2 (Monetary Reward: high, low) factorial design. We hypothesized that the positive expected utility associated with the main effect of reward will be expressed in the vmPFC and ventral striatum (Bartra et al., 2013; Levy & Glimcher, 2012; Sescousse et al., 2015), and the negative expected disutility of pain, most frequently associated with activations in the insula and anterior cingulate cortex (Atlas & Wager, 2012; Garcia-Larrea & Peyron, 2013), will be associated with the main effect of pain. Instead of using a region-of-interest approach, our design allowed us to functionally localize regions sensitive to utility (and disutility) by examining the main effects of pain and reward. Furthermore, we expected an interaction between pain and reward, such that in both sets of functionally-defined regions, the expected utility/disutility signal will be attenuated when a high magnitude of one outcome is accompanied by a high magnitude of an outcome with the opposite sign.

## Method

Participants undertook three calibration procedures in the following order: face rating task, pain scaling task, and reward scaling task. At the end of the three calibration procedures, the conditioning task began. The first block of the conditioning task was performed outside the scanner, and the next 5 blocks during fMRI scan. The conditioning task utilized 4 different compound USs, constructed from two individual levels of reward and stimulation. The two levels of reward were the high reward, determined in the reward scaling task (US_RH), and a low reward of 1p (US_RL). The two levels of stimulation, high (US_PH) and low (US_PL), were determined via the pain scaling procedure. The 4 face stimuli that acted as the CS in the conditioning task were selected from the face rating task and were randomly assigned to one of the four possible outcomes. CS_PHRH predicted high stimulation and high reward, CS_PHRL predicted high stimulation and low reward, CS_PLRH predicted low stimulation and high reward, and CS_PLRL predicted low stimulation and low reward.

### Participants

Twenty-five participants took part in the experiment. Technical problems necessitated removal of data from 3 participants, as well as data from the last 2 blocks (of 5) in one participant, and data from the last block in two others. Data from a further 2 participants was removed, one due to motion artefacts, and the other due to the participant’s failure to correctly demonstrate a consistent understanding of the CS-US contingencies during scanning (see Behavioural Results section). The final sample therefore included 20 participants (11 male, mean age 23.28, S.D = 4.14). The missing rating data for two participants from the final block was replaced by the linear trend at point. Likeability and threat ratings are reported from 19 participants, as the aforementioned technical problems prevent collection of the final ratings for one participant. The experiment received ethical approval from the Northwest 6 Research Ethics committee (Greater Manchester South).

### Apparatus

The electrical stimulations were delivered to the back of the right hand via an inhouse built ring electrode (Medical Physics, Salford Royal Hospital) attached to a Digitimer DS5 Isolated Bipolar Constant Current Stimulator (www.digitimer.com). To counter the effect of the magnetic field of the MR scanner, the DS5 stimulator was placed within a custom-built faraday cage (Medical Physics, Salford Royal Hospital). For reasons of participant safety, this stimulator was limited to delivering a maximum of 10mA during the experiment. Skin Conductance Response (SCR) was sampled at 600Hz using electrodes attached to a magnetically-sealed skin conductance machine (Medical Physics, Salford Royal Hospital). To ensure adequate conductance between the electrode and the skin, the back of each participant’s hand was prepared with Nuprep Skin Preparation Gel and Ten20 Conductive Paste. The SCR electrodes were placed on the inside medial phalange of the second and fourth fingers of the participant’s left hand. Both the outputs from the SCR machine, and the inputs to the DS5 machine were controlled via 1401plus data acquisition interface connected to a laptop running the Spike2 software (Cambridge Electronic Designs, Cambridge, UK). Both the laptop and the 1401 machine were kept inside the scan control room during data collection. The 1401 machine was connected to the skin conductance machine using a fibre-optic cable, and to the faraday cage using a BNC cable. To avoid interference between the faraday cage and the scanner, the faraday cage was kept a minimum of 2m from the scanner. The experiment was delivered via Cogent2000 on a Matlab platform (www.mathworks.com). Issues with the SCR data collection meant that many datapoints needed to be excluded from the subsequent analysis. As the analysis of the remaining data revealed no significant effects, this analysis is not discussed further.

### Materials

10 images of neutral male faces taken from Karolinska directed emotional faces database (Lundqvist et al., 1998) were used in the face rating task. Participants also filled out the following questionnaires: Spielberger State-trait Anxiety Inventory (Spielberger et al., 1983), BIS and BAS scales (Carver & White, 1994) and the Barratt Impulsivity Scale (Patton et al., 1985) as part of a wider study investigating the relationship between anxiety, impulsivity and the neural response to pain. Results relating to this questionnaire data are not therefore discussed further here.

### Procedure

Participants undertook three procedures in the following order: face rating task, pain scaling task, and reward scaling task. At the end of the three scaling procedures, the conditioning task began. The first block of the conditioning task was performed outside the scanner, and the next 5 blocks during fMRI scan.

#### Face rating task

This procedure ensured that the participant’s baseline attitude to the images used as the CS each condition of the conditioning task was similar. Participants were twice presented with 10 neutral faces in succession and were asked to rate how ‘threatening’ or ‘likeable’ they found each of the faces using a 9-point Likert scale. The presentation order of the faces, and the order in which the rating dimensions (likeability/threat) were applied, was random. Once the rating was complete the faces were ranked using an amalgamated like/threat score (threat ratings were first inverted, to get them onto the same scale as likeability scores), using the formula: 9-(threat score-1)+likeability score. The 4 median faces (i.e. those ranked 4, 5, 6 & 7) were selected for use as CS stimuli.

#### Pain scaling task

This procedure allowed us to identify two levels of electrical stimulation required for the conditioning task: a non-painful, but still noticeable stimulation level and a painful but still tolerable level (herein referred to as ‘low’ and ‘high’ stimulation). During the procedure, participants received a succession of stimulations which started with the input signal from the 1401plus interface to the DS5 set at 0.2V, and incremented at levels of 0.2V, up to a maximum of 5V. Participants rated each stimulation on a scale from 0 – 10 where a score of 0 reflected not being able to feel the stimulation, 4 reflected a stimulation that was on the threshold of being painful, 7 related to a stimulation that was deemed ‘painful but still tolerable’ and 10 related to ‘unbearable pain’. The scaling procedure was terminated once the participant reported the level of pain as being equivalent to ‘7’ on the scale. The procedure was performed twice, both times with a one-step-up method, to allow for initial habituation/sensitisation to the stimulation. The voltage levels rated as ‘1’ and ‘7’ on the second scaling were used for the ‘low’ (US_PL) and ‘high’ (US_PH) stimulation levels during the conditioning task.

#### Reward scaling task

This procedure was used to ensure that the appetitive US would have an equivalent subjective value to that of the aversive US (i.e. the high stimulation level). Participants were given the opportunity to make a monetary bid for the ‘pain’ stimulus using a Becker-deGrout-Marshak auction procedure (Becker et al., 1964). Each participant was asked to select a value equivalent to the minimum reward they were prepared to accept in order to experience a single high stimulation. A scale ranging from 5 to 50 British pence, incrementing in 5 pence steps, was used for this purpose. Participants were informed that the computer would also pick a value at random from the scale and that if the value they selected was lower or equal to the computer-generated value then they would receive both the money and the shock, but if the value they selected was higher than the computer-generated one, then they would receive neither. The value the participant chose was used as ‘high reward’ (US_RH) during the conditioning task. Once participants made their selection a “price” was selected at random, and determined whether participants received the high stimulation plus the high reward, or neither. This auction procedure ensured that participants did not just select the maximum value available. Participants were also given the option to indicate if they would require more than 50 British pennies in order to experience the pain stimulation, in which case the experiment was terminated, but no participant made that choice.

#### Conditioning task

Before the task started participants were informed of which outcome was associated with which CS, but not about the frequency with which each outcome would occur. Participants were told that they should keep track of the associations between the outcomes and the CS during the task, as after each block they would be required to estimate how often the ‘Pain’ stimulation and high reward occurred after each CS.

The trial design and timings were based on those used by (Bach et al., 2010). Each trial began with a 250ms fixation cross, this was then followed by the appearance of one of the CS images for 3500ms, shifted 50 pixels to the right or the left of the centre of the screen. To maintain engagement, participants were required to respond using the keypad to identify whether the image appeared to the left or right of the screen. Immediately after CS offset the US (combination of electrical stimulation and reward) either was or was not delivered. Each CS predicted its outcome occurring on 50% of trials. On trials where a US was delivered the participant received a 5ms electrical stimulation to the back of the hand while the monetary amount gained was shown in text (e.g. ’30p’ in the centre of the screen for 1000ms). On trials where the US was not delivered a 1000ms blank screen was shown. A jittered inter-trial interval (4s, 5s, 6s,7s or 8s) occurred before the start of the next trail. There were 5 experimental blocks, each conducted in a separate fMRI run. Each block contained 40 trials, 10 of each CS, of which 5 were reinforced. The presentation order of these trials was randomised within each block.

At the end of each block participants undertook 2 short tasks: pain experience ratings and US expectancy ratings. Firstly, they received one further (reinforced) presentation of each CS and were asked to rate their experience of the stimulation received on a scale from 0 (no sensation) to 100 (very painful). This task was included firstly to identify whether the concurrent delivery of different reward levels altered the experience of the stimulation, and secondly as a check for habituation or sensitisation to the stimulation. The second task involved the participants being presented with each CS in turn and being asked to estimate how often the CS had been followed by 1) the high pain stimulation and 2) the high reward level. This task was included to check that the participant was aware of the CS-US contingencies.

Participants undertook one training block of trials outside the scanner to ensure that they understood the CS-US contingencies before they went into the scanner.

Participants were deemed to be contingency-aware if they rated the ‘high pain’ and ‘high reward’ outcomes as occurring at least 20% of the time after the CSs which predicted these outcomes, and less than 20% of the time after the CSs which did not predict these outcomes. If the participant was unable to demonstrate this understanding on any of the CS’s, then they were then exposed to a second, shortened (24 trials with each CS) block of trials and their contingency understanding was rechecked. 7 participants received this one additional block of trials to demonstrate contingency awareness.

At the end of the training blocks outside the scanner and at the end of the scanning session participants completed 2 further tasks. In the first task participants were shown two of the CS side by side and were asked which they would prefer to see if they experienced another trial. There were three stages to this task, where the participant was asked to choose between the CS_PHRH and each of the other a CS in turn. The order in which the other 3 CS appeared was randomised. In the second task were also asked to complete ‘likeability’ and ‘threat’ ratings for each CS using the same scale used during the initial face rating task.

#### Image Acquisition

Participants were scanned using a 3T Philips Achieva scanner, fitted with a Philips 32-channel receive-only coil. Whole-brain functional images were collected using a single-shot dual-echo protocol (Halai et al., 2014), with TR=3s, TE=12ms & 35ms, FOV=240,240,132, flip angle=85. In each volume 33 slices of voxel size 3x3x4mm were collected in ascending order. Volumes were sampled at a 30 degree angle from the AC-PC line, as piloting revealed that this acquisition protocol produced the lowest signal drop out from the orbito-frontal cortex and amygdala. Prior to the functional scans, a whole-brain T1-weighted anatomical scan was acquired from each participant (TR=8.4s TE=3.8s, flip angle=8).

### Data analysis

#### Behavioural data analysis

Accuracy in the ‘left-right decision’ task relating to the CS were calculated for each of the 4 CS conditions and were entered into separate 2x2 ANOVAs with Conditioned Pain (high, low) and Conditioned Reward (high, low) as the factors. The pain experience ratings, pain expectancy, and reward expectancy, which were taken after each block, were entered into a 2x2x5 ANOVA. The Likeability and Threat ratings acquired at the end of the experiment were likewise entered into 2X2 ANOVAs. Violations of the assumption of sphericity during this analysis were corrected using the Greenhouse-Geisser correction.

#### MR data analysis

*Pre-processing.* MRI data was pre-processed and analysed using SPM12 (http://www.fil.ion.ucl.ac.uk/spm/). Prior to the analysis the first 5 volumes of each run were discarded to allow the scanner to reach magnetization equilibrium. Pre-processing began by manually centring the structural image to the Anterior Commissure using SPM’s ‘Set Origin’ function. Functional images were then realigned to the mean of the first run using a 6 parameter affine transformation and interpolated using a 4^th^ degree B-Spline. Slice time correction was then performed so that each slice was snapped to the 12^th^ slice within the volume (chosen to enhance signal from the ventral striatum, a region which has been associated with both appetitive and aversive prediction errors, Seymour et al., 2007). The individual’s structural scan was co-registered to the mean of these motion corrected functional images using a 12-parameter affine transformation. The structural scan was then segmented into separate images for white matter, grey matter and CRF and the resulting forward deformation field was used to normalise the functional data into MNI space. Finally the normalised images were smoothed using an 8mm FWHM isotropic Gaussian kernel.

##### Single subject models

Single subject data was modelled using the general linear model (GLM). Each experiment block was modelled using 9 regressors: each type of the non-reinforced CS, each type of US, and all reinforced CSs together. The reason for not separating reinforced CSs was that they were closely followed by the US. The regressors which coded CSs used a duration of 3.5 seconds as per the duration of CS presentation, and the others a duration of 0. Because it was possible that the hemodynamic response to the CS was associated with CS offset rather than onset, and because this could differ as a function of anticipating pain or reward, the BOLD response was modelled using a canonical HRF function alongside time and dispersion derivatives (Henson et al., 2001), improving overall model fit and thus residual variance assumption of the hierarchical modelling approach. For the same reason, CS regressors included parametric modulators that coded the latency to respond to each CS. In order to improve signal-to-noise ratio, nuisance regressor were entered into the model to account for the effect of motion, including the censoring of periods of extreme motion, using Siegel et al’s (Siegel et al., 2014) method implemented in the SPM utility plus toolbox (https://github.com/CPernet/spmup/blob/master/QA/spmup_first_level_qa.m). Six contrasts were computed at the single subject level and taken up to group models, corresponding to the canonical HRF for each of the non-reinforced CSs (CS_PHRH, CS_PHRL, CS_PLRH, CS_PLRL), as well as the main effect of pain [US_PH>US_PL] and main effect of reward [US_RH>US_RL].

##### Group models

For the key analysis of expected utility (response to CSs), the contrasts which corresponded to the canonical HRF response to non-reinforced CSs, were entered to a Conditioned pain (high, low) x Conditioned reward (high, low) flexible factorial design. The same model was used to examine effects of experienced utility (response to USs), with the same factors, but with contrasts corresponding to the canonical HRF response to USs. Each of the flexible factorial designs specified [dependence: yes, variance: unequal] for the subject factor, and [dependence: no, variance: equal] for other factors in the model.

##### Statistical inference

To analyse the pain and the reward models, we used an F identity contrast to describe significant effects of either regressor. These analyses used a cluster size FWE correction (Worsley et al., 1995) at *p*<.05. We applied a voxel discovery threshold of *p*<.0001 and a voxel extent threshold of k=10. We explored the main effects with t-contrasts (high>low reward, high>low pain), and the interaction with an F-contrast. We followed up on interaction effects by examining the four simple effects (Conditioned Pain under high reward, Conditioned Pain under low reward, Conditioned Reward under high pain, and Conditioned Reward under low pain). Analysis of simple effects used the same statistical thresholds as above, but only within a focused search volume, constrained to include only regions that expressed the interaction contrast at *p*<.05. Anatomical labelling used the neuromorphormetrix toolbox.

## Results

### Behavioural results

The behavioural data, shown in Figure 2, suggested that the manipulations were successful. The ratings taken after each block suggested that the effects were relatively consistent across the entire session.

**Figure 1.**
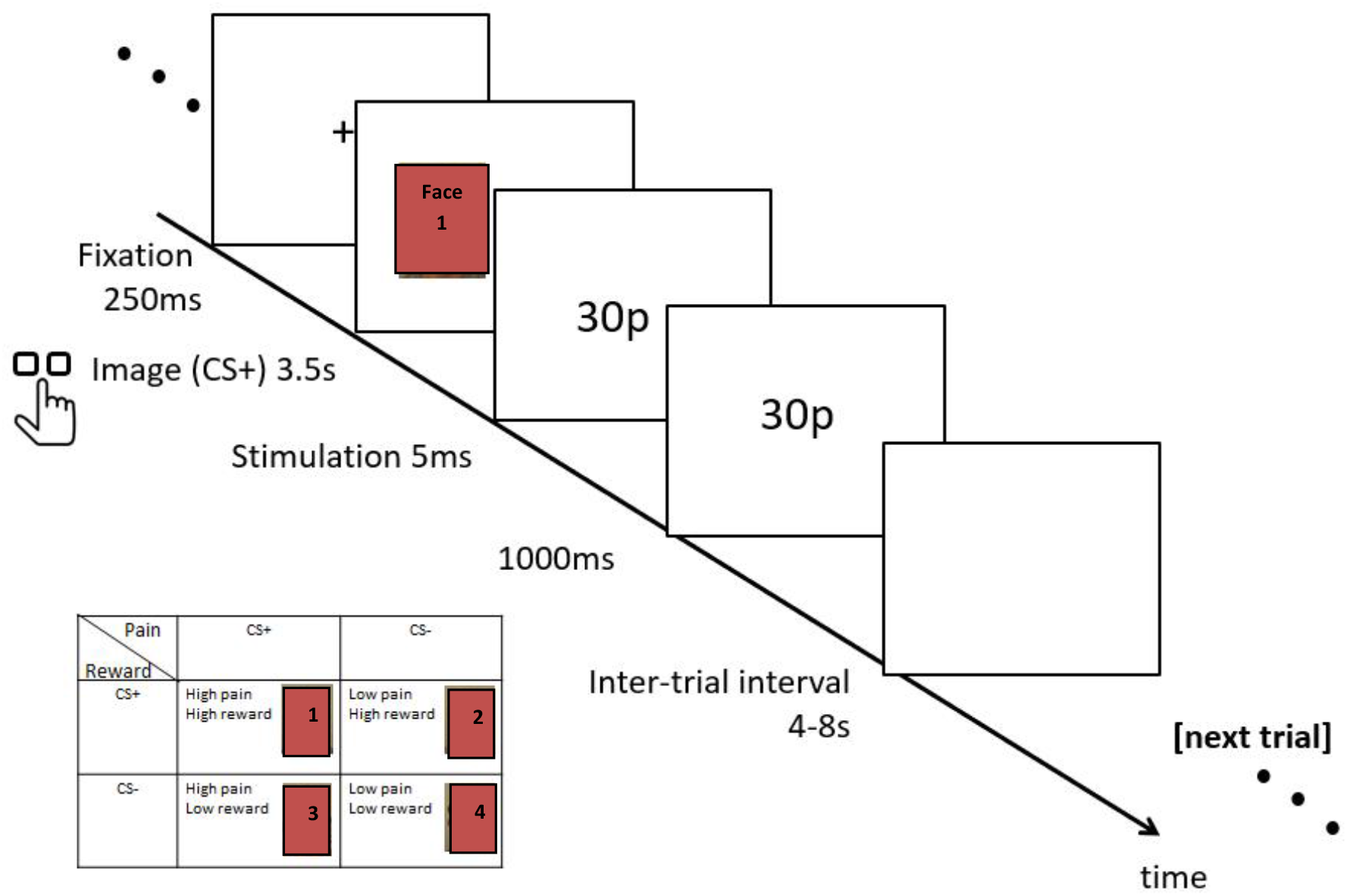
Experimental paradigm. Left insert: experimental design, showing that four faces were used in each of four conditions (P = pain, R = reward): CS_PHRH, CS_PHRL, CS_PLRH, CS_PLRL. The timeline a single trial with the CS_PHRH face is depicted. The dark squares in the pre-print cover the faces that were used in the study with the numbers 1… 4 denoting the 4 different faces.

**Figure 2:**
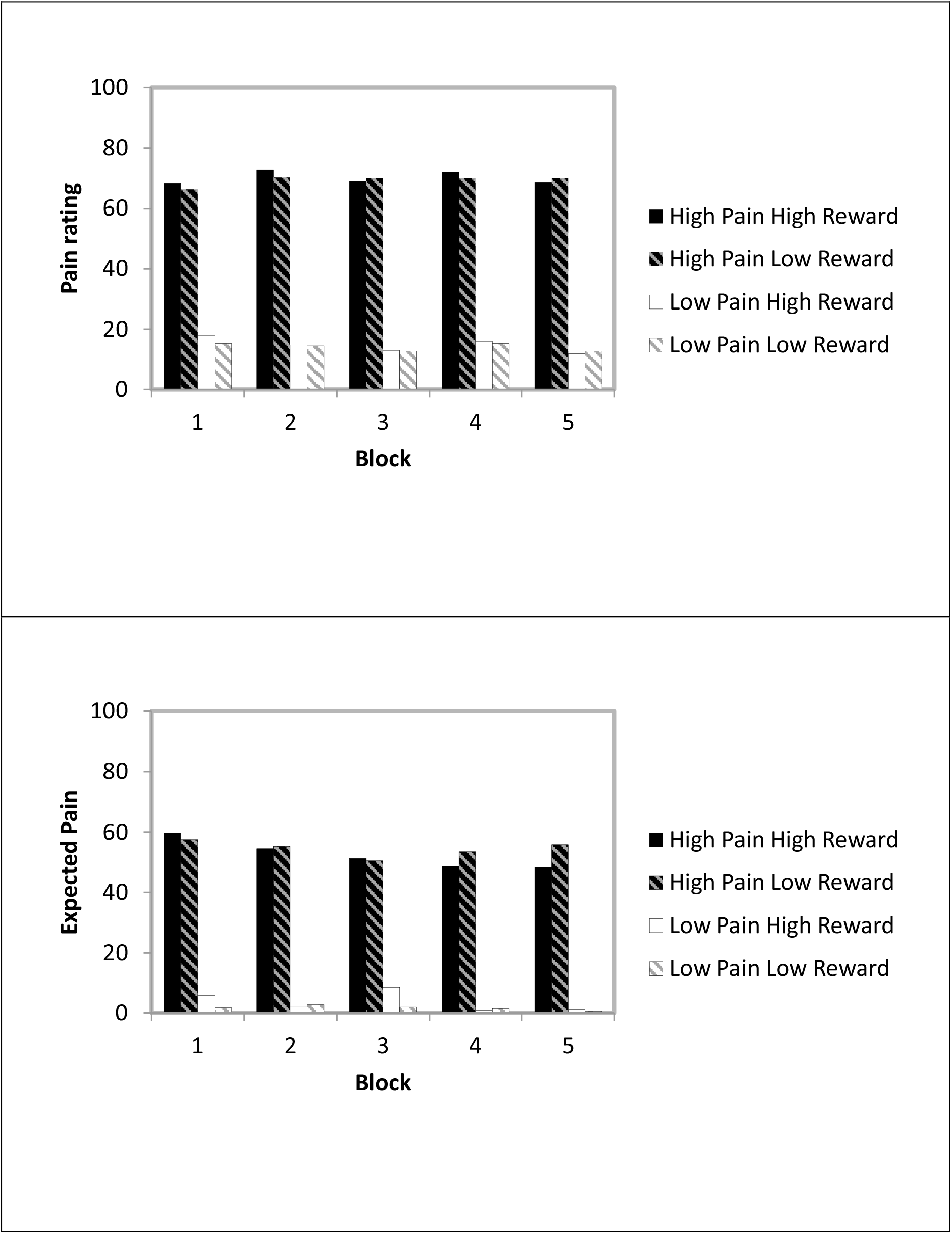
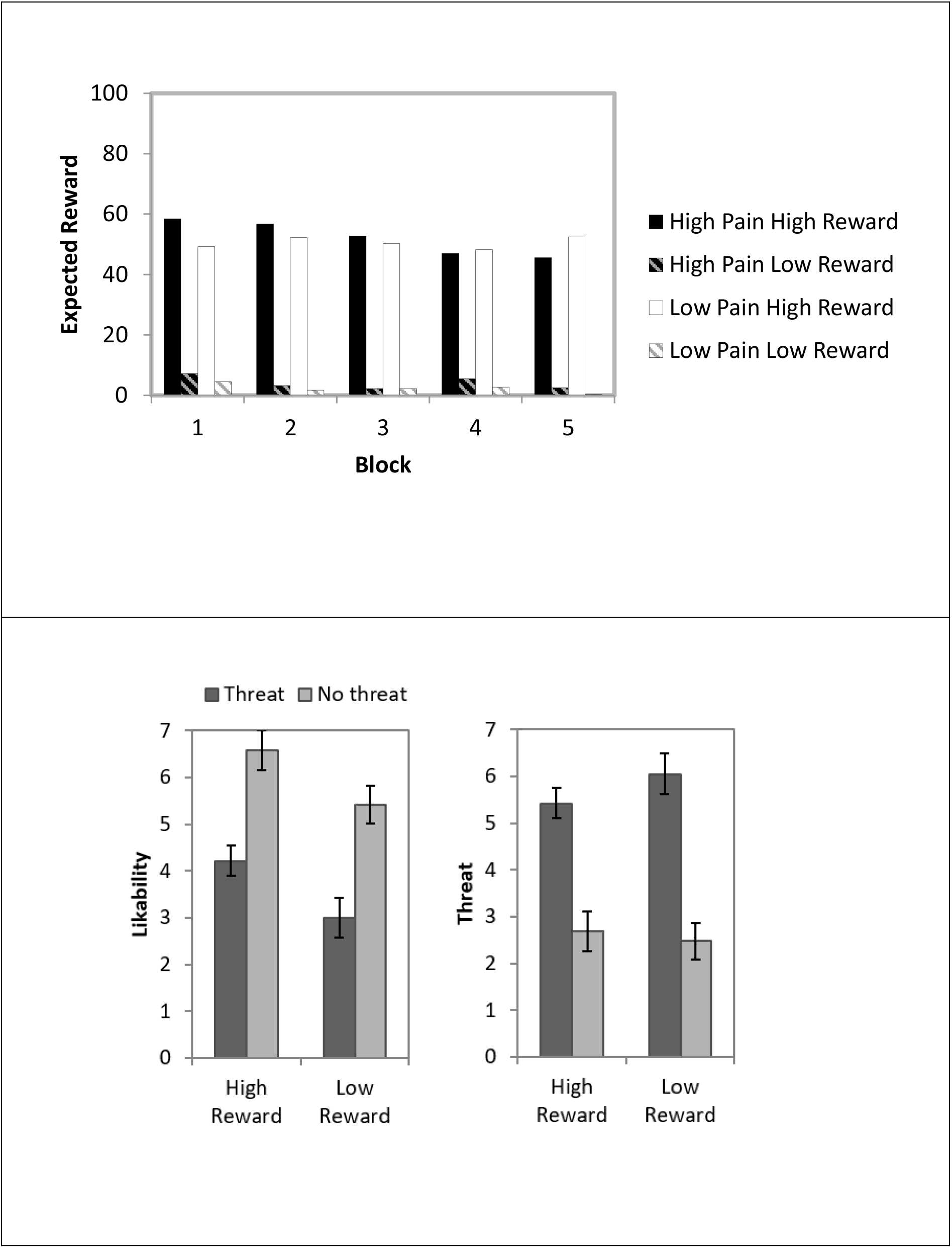
Behavioural data. The first three panels depict the average ratings of experienced pain, expected pain, and expected reward, based on ratings taken after each block. The bottom panel depict liking and threat ratings taken after the last block.

#### Task performance

During the CS task, conditioned pain decreased the accuracy by 3% to respond to the CS but conditioned reward did not (<1% change) and their interaction was not significant (3% vs 2%). Neither factors, nor their interaction, influenced the reaction times (755ms for CS_PHRH, 747ms for CS_PHRL, 758ms for CS_PLRH, 756ms for CS_PLRL – see table 1).

**Table 1:** Task performance statistics.

|  | Accuracy | Reaction Times |
| --- | --- | --- |
| Main effect pain | $F(1,19)=5.09$, $p=.036$, $\eta^2=.21$ | $F(1,19)=.29$, $p=.60$, $\eta^2=.01$ |
| Main effect reward | $F(1,19)=.03$, $p=.86$, $\eta^2=.00$ | $F(1,19)=.86$, $p=.37$, $\eta^2=.04$ |
| Interaction | $F(1,19)=.54$, $p=.47$, $\eta^2=.03$ | $F(1,19)=.08$, $p=.78$, $\eta^2=.00$ |

#### Ratings of pain experience

The analysis of the pain ratings given at the end of each block in response to each US, revealed that the high pain US was experienced as more painful that the low pain US (Table 2). Importantly there was no interaction with block, such that high pain was rated, on average, as 67 after block 1 and 69 after block 5, and the difference between high and low pain 51 points after block 1 and 57 points after block 5, suggesting that there was not significant habituation to the pain stimuli. There was no impact of reward on the pain experience ratings.

**Table 2:** Statistics for ratings of pain experience, pain expectancy and reward expectancy.

|  | Pain experience | Pain expectancy | Reward expectancy |
| --- | --- | --- | --- |
| Main effect pain | $F(1,19)=214.05$, $p=.001$, $\eta^2=.92$ | $F(1,19)=287.14$, $p<.001$, $\eta^2=.94$ | $F(1,19)=2.09$, $p=.16$, $\eta^2=.1$ |
| Main effect reward | $F(1,19)=.45$, $p=.51$, $\eta^2=.02$ | $F(1,19)=0$, $p=1$, $\eta^2=.0$ | $F(1,19)=227.15$, $p<.001$, $\eta^2=.91$ |
| Interaction pain x reward | $F(1,19)=.04$, $p=.84$, $\eta^2=0$ | $F(1,19)=3.87$, $p=.06$, $\eta^2=.17$ | $F(1,19)=.04$, $p=.84$, $\eta^2=.00$ |
| Main effect block | $F(4,76)=1.42$, $p=.24$, $\eta^2=.07$ | $F(4,76)=2.70$, $p=.04$, $\eta^2=.12$ | $F(4,76)=.1.77$, $p=.14$, $\eta^2=.08$ |
| Interaction pain x block | $F(4,76)=1.68$, $p=.20$, $\eta^2=.08$ | $F(4,76)=1.83$, $p=.13$, $\eta^2=.14$ | $F(2.53,48.02)=1.46$, $p=.24$, $\eta^2=.07$ |
| Interaction reward x block | $F(4,76)=.72$, $p=.52$, $\eta^2=.04$ | $F(2.48,47.09)=3.13$, $p=.043$, $\eta^2=.14$ | $F(2.26,43.04)=.94$, $p=.47$, $\eta^2=.05$ |
| Interaction pain x reward<br>x block | $F(4,76)=.21, p=.83,$<br>$\eta^2=.01$ | $F(4,76)=2.68,$<br>$p=.07, \eta^2=.12$ | $F(2.43,46.26)=1.08, p=.37,$<br>$\eta^2=.05$ |

#### Ratings of US expectancy

The participant’s expectancy ratings indicated how often they expected that each CS was followed by the pain and reward USs. Their ratings confirmed that they consistently understood the predictive characteristics of each CS (although note that one participant was excluded entirely from the analysis for failing to demonstrate this). Participants expected the high pain US more often after the CS_PHRH and CS_PHRL than the other stimuli. Interestingly the effect of block on pain expectancy rating was significant, and interacted with conditioned reward, such that participants expected high pain less frequently in later blocks, with this reduction more pronounced when the CS also predicted reward: participants thought the high pain occurred more frequently under the CS_PHRL than the CS_PHRH as the task went on (Figure 2). The other main effects and interactions were not significant (Table 2). The participants also demonstrated understanding of the reward contingencies, in that they expected reward more often after the CS_PHRH and CS_PLRH than the other stimuli. The other main effects and interactions were not significant (Table 2).

#### Ratings of CS Likeability, threat and preference

*C*onditioned pain decreased face likeability ratings by 2.4 points, and conditioned reward increased them by 1.18 points. The interaction was not significant (2.37 vs. 2.42 points; Table 3). Conditioned pain rendered faces more threatening by 3.16 points, but conditioned reward did not (0.2 points decrease); the interaction was not significant (2.74 vs. 3.58 points; Table 3).

**Table 3:** Statistics for likability and threat ratings.

|  | Likability | Threat |
| --- | --- | --- |
| Main effect pain | $F(1,18)=19.75, p<.001, \eta^2=.52$ | $F(1,18)=33.43, p<.001, \eta^2=.65$ |
| Main effect reward | $F(1,18)=10.75, p=.004, \eta^2=.37$ | $F(1,19)=.81, p=.38, \eta^2=.04.$ |
| Interaction | $F(1,18)=0, p=.94, \eta^2=.00.$ | $F(1,18)=1.83, p=.19, \eta^2=.09$ |

At the end of the experiment, the majority (14 of 19) participants chose the CS_PHRH over the CS_PHRL, and 11 (out of 19) chose it over the CS_PLRL, suggesting that the preferred to have the reward even at the cost of pain.

### fMRI results

#### Main effects of pain and reward USs

The main effect of US pain resulted in a robust activation in the primary and secondary somatosensory cortices, postcentral gyrus, and right anterior insula (Figure 3 and Table 4). Neither the response to reward nor the interaction of pain and reward survived our statistical threshold.

**Figure 3.**
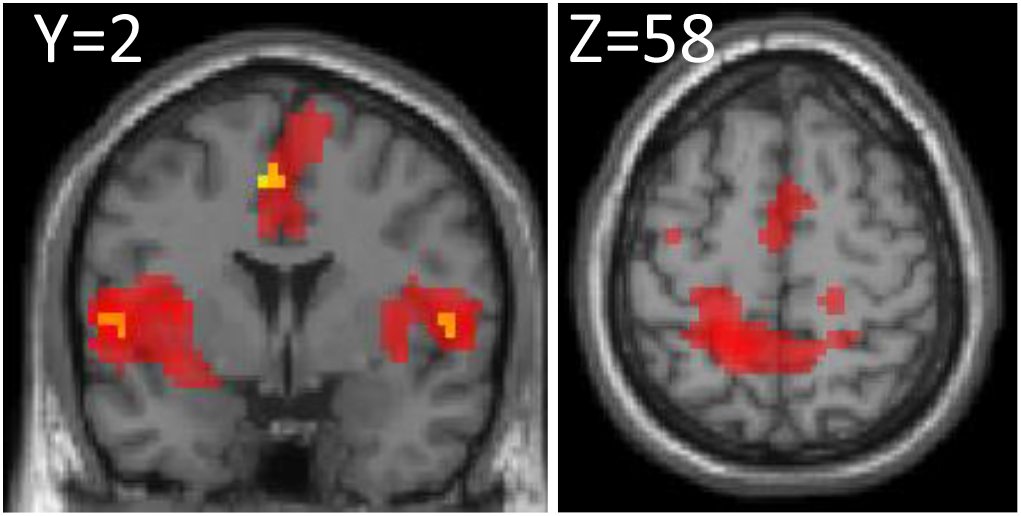
Main effects of experienced pain (red) and expected pain (yellow), cluster FWE<.05.

**Table 4.**
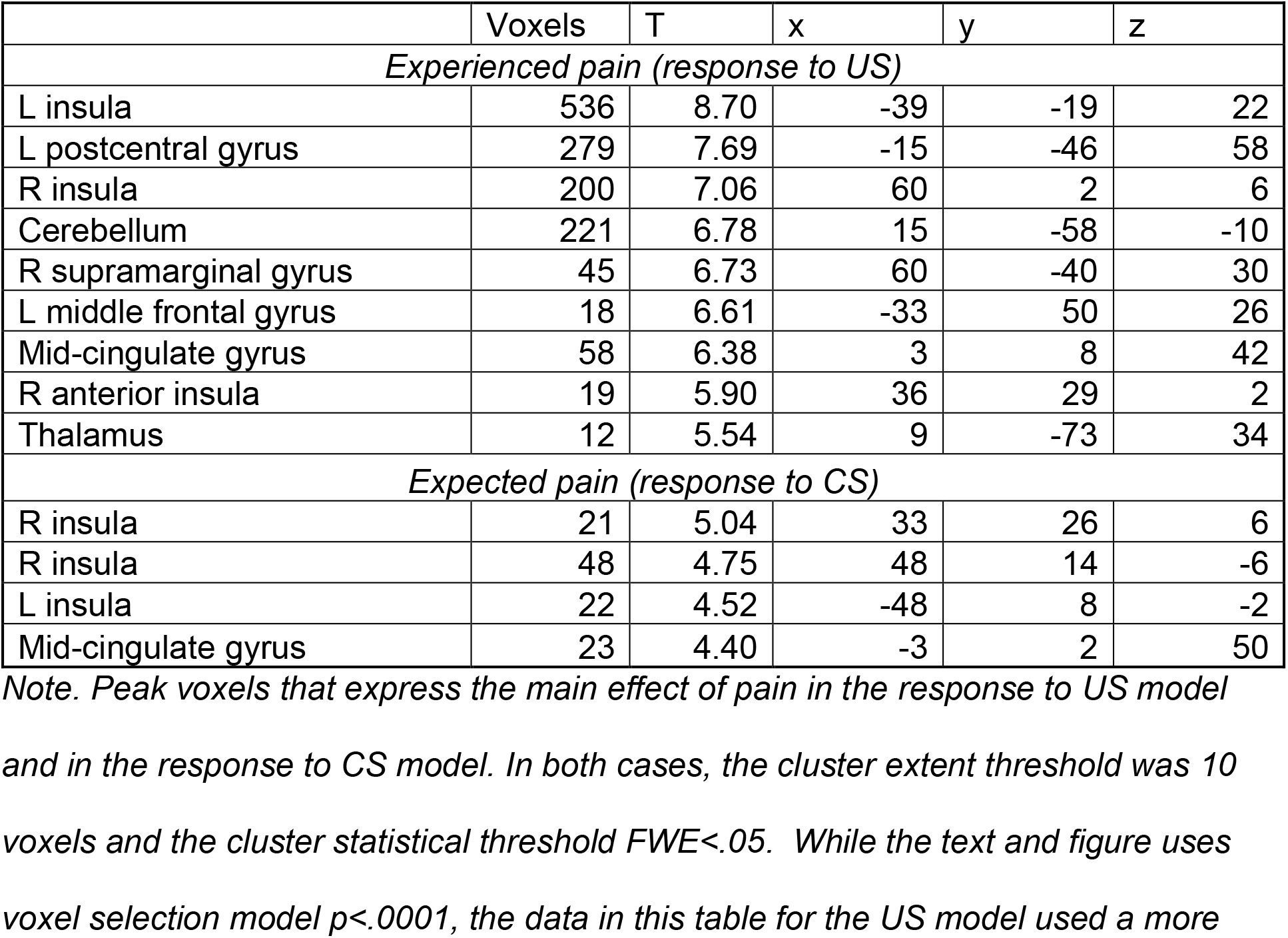
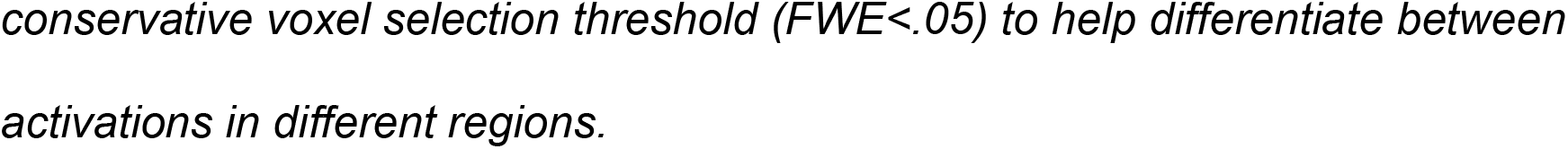
The main effect of pain and expected pain.

|  | Voxels | T | x | y | z |
| --- | --- | --- | --- | --- | --- |
| *Experienced pain (response to US)* |  |  |  |  |  |
| L insula | 536 | 8.70 | -39 | -19 | 22 |
| L postcentral gyrus | 279 | 7.69 | -15 | -46 | 58 |
| R insula | 200 | 7.06 | 60 | 2 | 6 |
| Cerebellum | 221 | 6.78 | 15 | -58 | -10 |
| R supramarginal gyrus | 45 | 6.73 | 60 | -40 | 30 |
| L middle frontal gyrus | 18 | 6.61 | -33 | 50 | 26 |
| Mid-cingulate gyrus | 58 | 6.38 | 3 | 8 | 42 |
| R anterior insula | 19 | 5.90 | 36 | 29 | 2 |
| Thalamus | 12 | 5.54 | 9 | -73 | 34 |
| *Expected pain (response to CS)* |  |  |  |  |  |
| R insula | 21 | 5.04 | 33 | 26 | 6 |
| R insula | 48 | 4.75 | 48 | 14 | -6 |
| L insula | 22 | 4.52 | -48 | 8 | -2 |
| Mid-cingulate gyrus | 23 | 4.40 | -3 | 2 | 50 |

*Note. Peak voxels that express the main effect of pain in the response to US model and in the response to CS model. In both cases, the cluster extent threshold was 10 voxels and the cluster statistical threshold FWE<.05. While the text and figure uses voxel selection model $p<.0001$ , the data in this table for the US model used a more*
conservative voxel selection threshold ( $FWE < .05$ ) to help differentiate between activations in different regions.

#### Effects associated with CSs

No voxel survived in the analysis of the main effect of Conditioned Reward. The main effect of conditioned pain activated the mid-cingulate and the insula bilaterally (Figure 3 and Table 4). Each of the peak voxels in the regions that responded to the main effect of conditioned pain was also significantly activated by the interaction contrast between pain and reward. We therefore decided to focus our analysis on the simple effects, which were masked by the interaction contrast, as described above. These are depicted in Figure 4 and Table 5.

**Figure 4.**
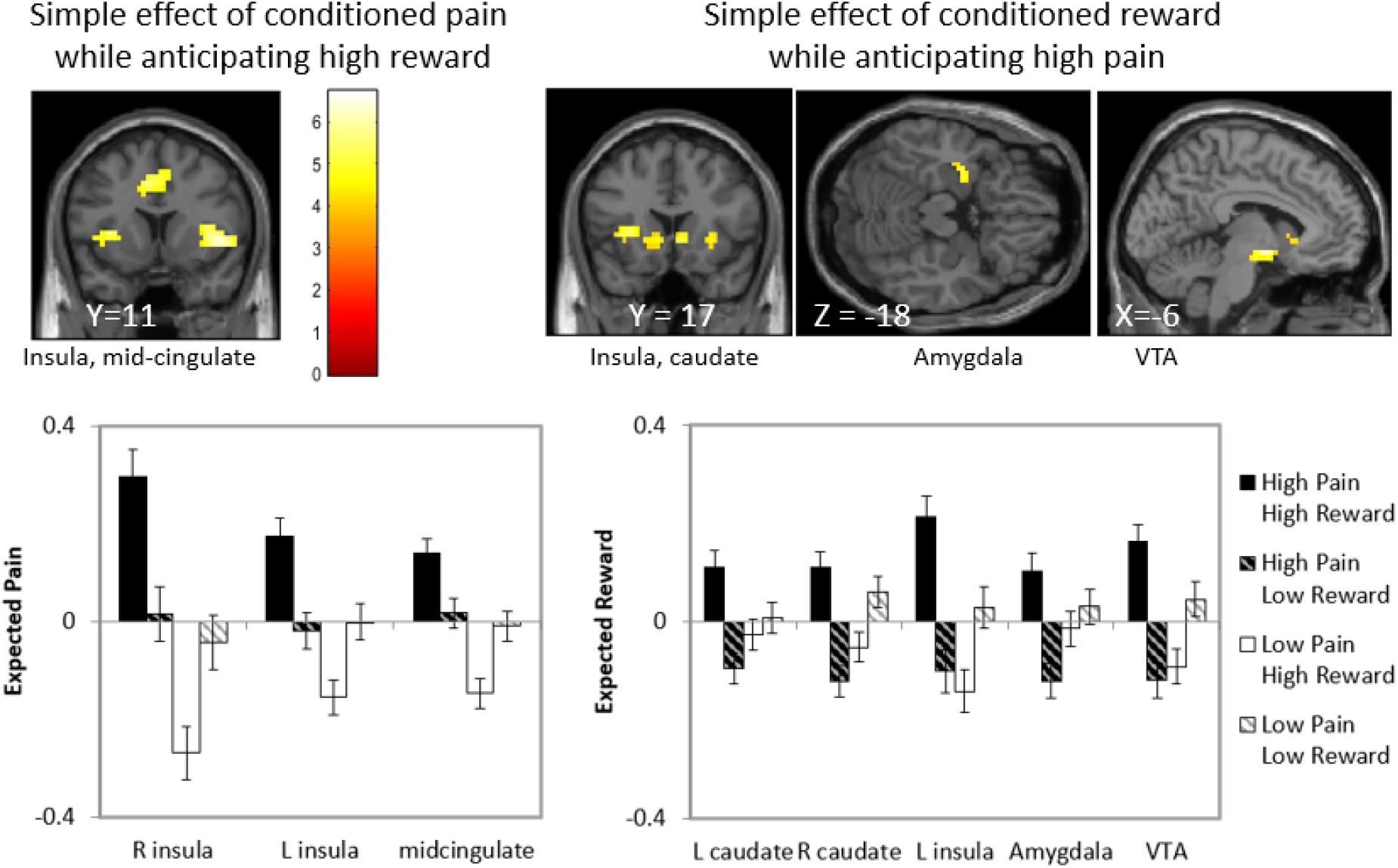
Simple effects of conditioned pain and conditioned reward, cluster threshold FWE<.05, masked by their interaction. LEFT: the simple effect of conditioned pain while participants were expecting high reward [CS_PHRH> CS_PLRH]. RIGHT: the simple effect of conditioned reward while participants were expecting high pain [CS_PHRH> CS_PHRL].

**Table 5.** Analysis of simple effects that unpacks the significant interaction between Conditioned pain and Conditioned reward.

|  | voxels | T | x | y | z |
| --- | --- | --- | --- | --- | --- |
| *Reward under high pain* |  |  |  |  |  |
| Ventral striatum | 91 | 6.72 | -9 | -1 | -10 |
| Cerebellum | 45 | 6.20 | 9 | -46 | -26 |
| L caudate extending to L insula and L amygdala | 159 | 6.13 | -18 | 26 | -10 |
| R caudate | 48 | 5.17 | 9 | 17 | -2 |
| R middle temporal gyrus | 19 | 4.59 | 48 | -4 | -26 |
| *Pain under high reward* |  |  |  |  |  |
| R insula/frontal operculum | 249 | 6.16 | 45 | 11 | 2 |
| R calcarine cortex | 70 | 6.00 | 18 | -76 | 6 |
| L insula/frontal operculum | 122 | 5.91 | -27 | 20 | 6 |
| Cerebellum | 36 | 5.91 | 9 | -52 | -26 |
| mid-cingulate gyrus | 146 | 5.56 | -6 | 5 | 46 |
| L precentral gyrus | 19 | 5.43 | -48 | -4 | 38 |
| L Superior parietal lobule | 88 | 5.36 | -36 | -43 | 66 |
| L calcarine cortex | 19 | 5.05 | -18 | -76 | 6 |
| Posterior cingulate cortex | 22 | 4.95 | -18 | -28 | 38 |
| R superior temporal gyrus | 20 | 4.84 | 60 | -37 | 14 |
| R cuneus | 22 | 4.78 | 3 | -79 | 22 |
| *Reward under low pain* |  |  |  |  |  |
| L superior frontal medial gyrus | 96 | 5.36 | -6 | 44 | 34 |
| R superior frontal medial gyrus | 20 | 4.87 | 21 | 41 | 38 |

Note. Peak voxels are presented in each of the simple effects that follow from the analysis of the interaction of pain and reward. They are masked inclusively by the $F$ interaction contrast, $p < .05$ , voxel extent = 10. Within the mask, the voxel selection threshold was $p < .0001$ , the cluster extent threshold was 10 voxels and the cluster statistical threshold $FWE < .05$ .

We found that the effect of conditioned pain was expressed in the mid-cingulate and bilateral insula when participants also expected high reward, but not when they expected low reward. In fact, the simple effect of conditioned pain under low reward did not result in any significant activations. The simple effect of reward under high pain activated the VTA, and the caudate nucleus and insula bilaterally. The simple effect of reward under low pain activated the left angular gyrus and the left superior frontal gyrus.

## Discussion

We have manipulated reward and pain using the same Pavlovian conditioning procedure where neutral faces predicted combinations of pain and reward to examine how the brain represents a stimulus that predicts a mix of positive and negative outcomes. The behavioural data suggested that participants understood the contingencies between CSs and USs, had a consistent experience of pain throughout the experiment, and, crucially, their threat and likeability ratings suggested that they have acquired a conditioned response. Our neuroimaging results differed from those hypothesised based on previous studies (Bulganin et al., 2014; Choi et al., 2013; Yee et al., 2021), which reported attenuation of the neural markers of anticipation of the positive outcome (reward) in the presence of the negative outcome (pain), and vice versa. Instead, we observed enhanced expression of reward anticipation in the presence of high pain (compared to low reward), and enhanced anticipation of pain in the presence of high reward (compared to low reward).

Our results contradict a straightforward interpretation based on the subjective utility of the outcomes. Intuitively, the overall utility of reward would be lower when it is accompanied by high pain, and therefore a brain region that represents the simple effect of reward (high > low) should be less active under high than low pain. Likewise, the disutility of pain should decrease (get closer to zero) when accompanied by high reward, and therefore the simple effect of conditioned pain (high > low) should decrease under high than low reward. Therefore, although theories such as prospect theory (Kahneman & Tversky, 1979) have not been designed to handle multi-attribute prospects, it is reasonable to deduce that the pattern of activations we observed is unlikely to be expressing subjective utility. Although it is possible that the regions we observed in the analysis of simple effect were sensitive to expected sensory-specific aspects of the disutility of pain or their negative valence (Chikazoe et al., 2014), it is more difficult to argue for sensory-specific aspects of monetary rewards. Another framework that predicts that activation of the appetitive system by reward anticipation should attenuate the activation of the system that represents pain is the theory that motivational systems associated with approach (towards appetitive stimuli) and avoidance (of aversive stimuli) are mutually inhibitory (Konorski, 1967; Dickinson and Dearing 1979). The pattern observed here, however, aligns better with a facilitative effect of one system on another.

The conditioned stimulus that predicted a combination of high-pain and high reward was associated, numerically, with the greatest BOLD signal than all other stimuli. This was, arguably, the most salient of the four stimuli, and it is therefore reasonable to ask whether the pattern we observed in response to conditioned stimuli was due to salience. The aspect of salience most relevant here is motivational salience, defined by Schultz (Schultz, 2016) as “The ability of a stimulus to elicit attention due to its positive (reward) or negative (punishment) motivational value. Motivational salience is common to reward and punishment”. According to this definition, the conditioned stimulus that predicted a both high reward and high punishment would be more salient than conditioned stimuli that predicts just one of these outcomes, or lower levels of reward and punishment. Motivational salience is closely associated with emotional arousal; and both activate the insula and mid-cingulate (Lindquist et al., 2015; Seeley, 2019), which here were more activated for the CS predicting two motivationally-salient outcomes (high-pain, high-reward) compared to the CS predicted low-pain, high-reward. This interpretation is challenged by the finding that a different set of regions were more activated for the high-pain, high-reward CS when it was compared to the high-pain, low-reward CS, a comparison which should operationalise salience just as well, because in both cases we compare a stimulus that predicts two personally significant outcomes to a stimulus which predicts just one. Perhaps the asymmetry occurred because one of these two outcomes was less personally-significant than the other, despite our best efforts to equate their hedonic value. However, if one of the outcomes had very limited motivational impact, for example if participants did not care much for the monetary reward they could earn, or completely habituated to pain, we should have observed a strong main effect of the other, rather than a strong interaction. In addition, the pain ratings taken at the end of each block suggested that the experience of the high pain stimulus did not decrease significantly during the experiment.

One way to explain the pattern of findings observed here is by considering dynamic changes to participants’ reference point. In prospect theory, the reference point is fixed at zero gains and zero losses. There is evidence, however, that the reference point can be set endogenously, and that it often relates to the most salient aspects of a person’s local context (Kıbrıs et al., 2023). For example, consumers who invested in a particular product through a loyalty scheme, e.g. purchasing a particular brand for loyalty points, later preferred small sure rewards over larger uncertain rewards. Analysis showed that their invested effort was best modelled as having moved the reference point, so that the loyal consumers were essentially experiencing a deficit, and rewards were no longer (above-zero) gains moved them closer to zero utility (Kivetz, 2003). Because in prospect theory the utility function is steeper at the loss than the gain domain, having invested effort consumers were now more sensitive to smaller rewards than those who have not invested.

The same may have occurred in our experiment, although we offer this interpretation with utmost caution. Predicting high pain could shift the reference point to the loss domain, rendering participants more sensitive to expected reward. Such shift in reference point could explain why the difference between high and low reward in the caudate, insula, amygdala and VTA was greater when participants predicted high pain compared to low pain. This interpretation is supported by previous finding that when participants performed an instrumental task to gain reward that was accompanied by pain, pain increased sensitivity to reward (Talmi et al., 2009). It is possible that predicting high reward had similar effects – shifting the reference point so that participants became more sensitive to expected pain. This could explain the greater difference between the high and low pain in the insula and cingulate in the high reward condition, compared to the low reward condition. Figure 5 illustrates these shifts in reference points and how they could operate to increase sensitivity to expected rewards and expected pain. The figure also reveals that such effects would depend very much on the subjective utility of the outcomes and characteristics of the utility function. Therefore, the interpretation we offer here could be tested in future by varying the amounts of pain and reward parametrically.

**Figure 5.**
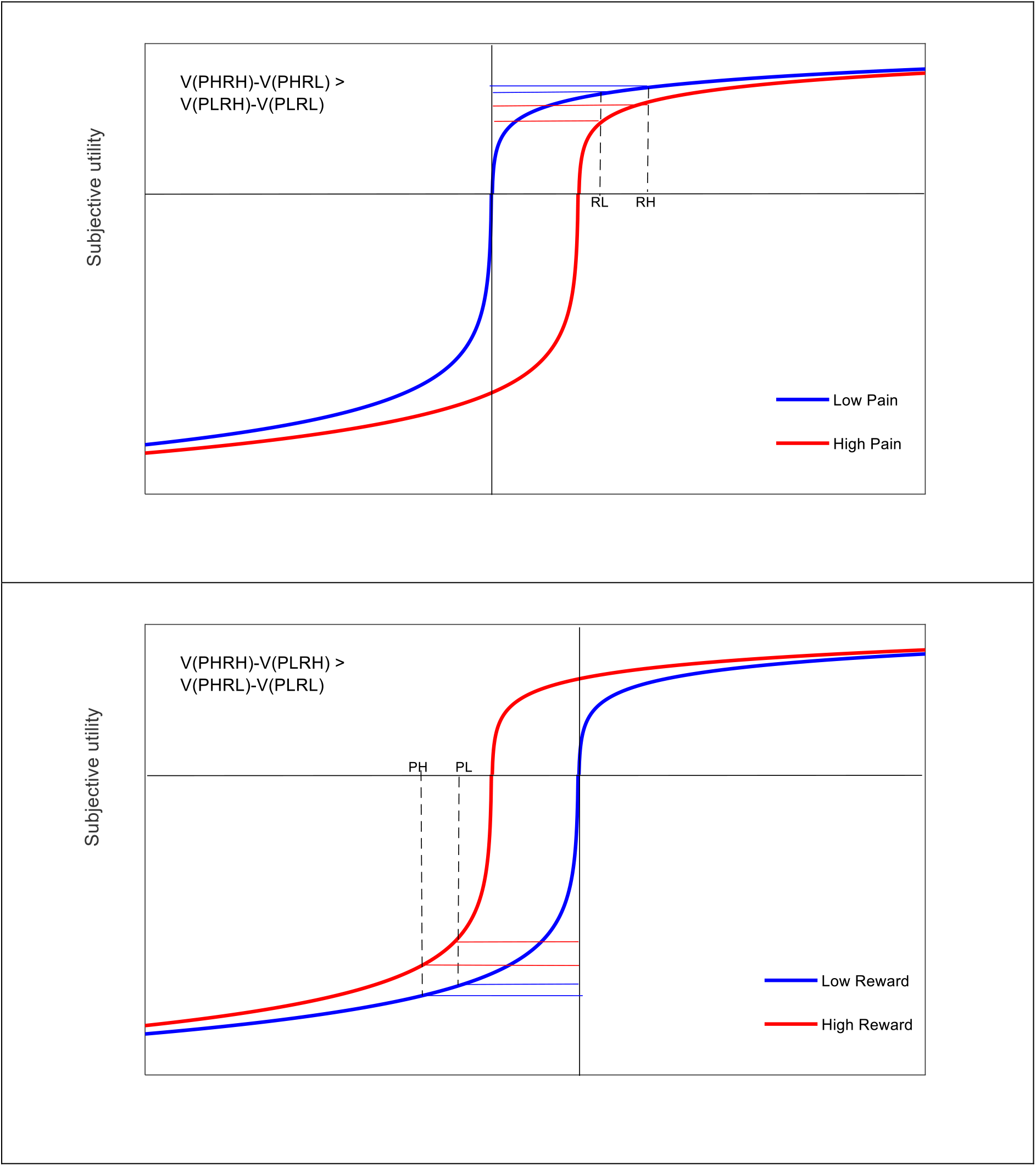
Changes in subjective utility expected on CS onset due to dynamic shifts in reference point. Conditions are labelled PHRH (high pain, high reward), PLRH (low pain, high reward), PHRL (high pain, low reward), PLRL (low pain, low reward). Top panel. Predicting high pain shifts the reference point rightwards (red curve) compared to when predicting low pain, so that the absence of reward (the y axis intercept) is experienced as an expected loss of utility. The subjective utility of monetary reward in the high pain condition, illustrated in the difference between high and low reward, is decreased compared the low pain condition (blue curve). Bottom panel. Predicting high reward shifts the reference point leftward (red curve) compared to when predicting low reward, so that the absence of pain (the y axis intercept) is experienced as an expected gain in utility. The subjective disutility of pain in the high reward condition, illustrated in the difference between high and low pain, is decreased compared the low reward condition (blue curve).

All economic models of individual choice include some form of gain-loss value integration, where multi-attribute prospects are only selected when their positive attributes (e.g. luxurious cocktails and brilliant conversation) compensate sufficiently for its negative attributes (e.g. exorbitant price and time costs). Our results contribute to understanding of what happens when incommensurate, opposite-valence attributes are represented simultaneously. They suggest that it is possible for people to become more sensitive to potential rewards when they consider that they may suffer a cost, and more sensitive to potential pain when there is also a chance to gain something at the same time.

## Acknowledgements

We thank AKP Jones and the Human Pain group, and S. Watson at the clinical engineering team, both at Salford Royal hospital, for their support. Funding: This work was supported by an ESRC First Grant [ESRC ES/I010424/1].

